# Logical modeling of dendritic cells *in vitro* differentiation from human monocytes unravels novel transcriptional regulatory interactions

**DOI:** 10.1101/2020.08.14.251710

**Authors:** Karen J. Nuñez-Reza, Aurélien Naldi, Arantza Sanchéz-Jiménez, Ana V. Leon-Apodaca, M. Angélica Santana, Morgane Thomas-Chollier, Denis Thieffry, Alejandra Medina-Rivera

**Affiliations:** Laboratorio Internacional de Investigación sobre el Genoma Humano, Universidad Nacional Autónoma de México, Juriquilla, México; Computational Systems Biology team, Institut de Biologie de l’Ecole normale supérieure, Inserm, CNRS, Université PSL, Paris, France; Centro de Investigación en Dinámica Celular (IICBA), Universidad Autónoma del Estado de Morelos, Cuernavaca, México

**Keywords:** Dendritic cells, differentiation, Logical modeling, Regulatory networks

## Abstract

Dendritic cells are the major specialized antigen-presenting cells, thereby connecting innate and adaptive immunity. Because of their role in establishing adaptive immunity, they have been used as targets for immunotherapy. Monocytes can differentiate into dendritic cells *in vitro* in the presence of colony-stimulating factor 2 (CSF2) and interleukin 4 (IL4), activating four signalling pathways (MAPK, JAK/STAT, NFKB, and PI3K). However, the transcriptional regulation responsible for dendritic cell differentiation from monocytes (moDCs) remains unknown. By curating scientific literature on moDCs differentiation, we established a preliminary logical model that helped us identify missing information for the activation of genes responsible for this differentiation, including missing targets for key transcription factors (TFs). Using ChIP-seq and RNA-seq data from the Blueprint consortium, we defined active and inactive promoters, together with differentially expressed genes in monocytes, moDCs, and macrophages (which correspond to an alternative cell fate). We then used this functional genomic information to predict novel targets for the identified TFs. We established a second logical model integrating this information, which enabled us to recapitulate the main established facts regarding moDCs differentiation. Prospectively, the resulting model should be useful to develop novel immunotherapies based on moDCs regulatory network.

## Introduction

Dendritic cells (DCs) are the main antigen-presenting cells (1), whose role is to activate the innate immune response, by presenting antigens to the naïve lymphocytes in order to initiate the immune response (2). Dendritic cells have been used in immunotherapies for their capacity to activate the adaptive immune response, in particular, dendritic cells derived from monocytes (moDCs) (3), as monocytes circulate in peripheral blood, they are easily accessible. Furthermore, there is an established protocol for moDCs differentiation (3).

The protocol to differentiate monocytes to moDCs consists in cultivating monocytes with colony-stimulating factor 2 (CSF2) and interleukin 4 (IL4) (4). When only IL-4 is used, monocytes are activated, while treatment with CSF2 results in their differentiation into macrophages. Only the combined stimuli results in DC differentiation, pointing to the importance of signalling interplay for the differentiation of moDCs. CSF2 signalling leads to the activation of NFΚB, MAPK, PI3K, JAK2, and STAT5 (5,6). IL4 signalling activates the JAK/STAT pathway, while JAK1 activates STAT3 and JAK3 activates STAT6 (7). There are some well-known transcription factors (TFs) ultimately activated by CSF2 and IL4 signalling pathways, but presumably, only a fraction of the target genes participating in moDCs differentiation have been reported (6,8,9).

A good way to integrate multiple signalling pathways into a comprehensive regulatory network and check its coherence consists of developing a dynamical model (10). As most of the available data are qualitative, it is natural to use a qualitative approach to build such a model. Logical models are well suited to represent this qualitative data and have been proposed for various similar processes (11–13). This qualitative formalism relies on the construction of a regulatory graph, whose nodes denote molecular components, while arcs denote (positive, negative, or dual) regulatory interactions. In the simplest cases, nodes are associated with Boolean variables, which take the values 0 or 1, denoting absence/inactivity or presence/activation, respectively (14). Logical models are usually derived based on a careful manual curation of relevant scientific literature; but they can also be enriched using other sources of information, such as high-throughput sequencing data (15). Logical models can integrate different kinds of molecular entities (genes, proteins, lncRNA, etc.) (15).

GINsim is a computational tool dedicated to the building and analysis of logical models, enabling the delineation of logical regulatory graphs, together with various dynamical analyses, through model simulations, but also with the support of efficient algorithms to identify the attractors (stable states and/or oscillatory behavior) of the system, for wild-type or mutant conditions (14). The resulting model can be further analysed using the CoLoMoTo tool suite, an interactive toolbox integrating several logical modeling software tools, with a uniform interface to perform simulations and other analyses, which are easy to share, and reproduce through the use of notebooks (16).

The aim of our study was to integrate all the information gathered from scientific literature and high-throughput data (RNA-seq and ChIP-seq) into a logical model of the regulatory network underlying moDCs differentiation. After iterative enhancement, our final model is able to properly recapitulate cell commitment for each of the initial conditions considered: (i) IL4 alone fosters monocyte activation, (ii) CSF2 alone fosters macrophage commitments, while (iii) CSF2 and IL4 together foster moDC commitment.

## Results

### Information gathered from literature curation leads to a fragmentary model of monocytes to dendritic cells differentiation

To better understand the regulatory network controlling moDCs differentiation, we analysed the scientific literature and integrated relevant information into a regulatory graph. In this process, we focused on monocyte to moDCs differentiation studies carried on human cells, in particular, on studies where CSF2 and IL4 were used in similar culture conditions. The resulting regulatory graph is shown in Figure 1.

**Figure 1.**
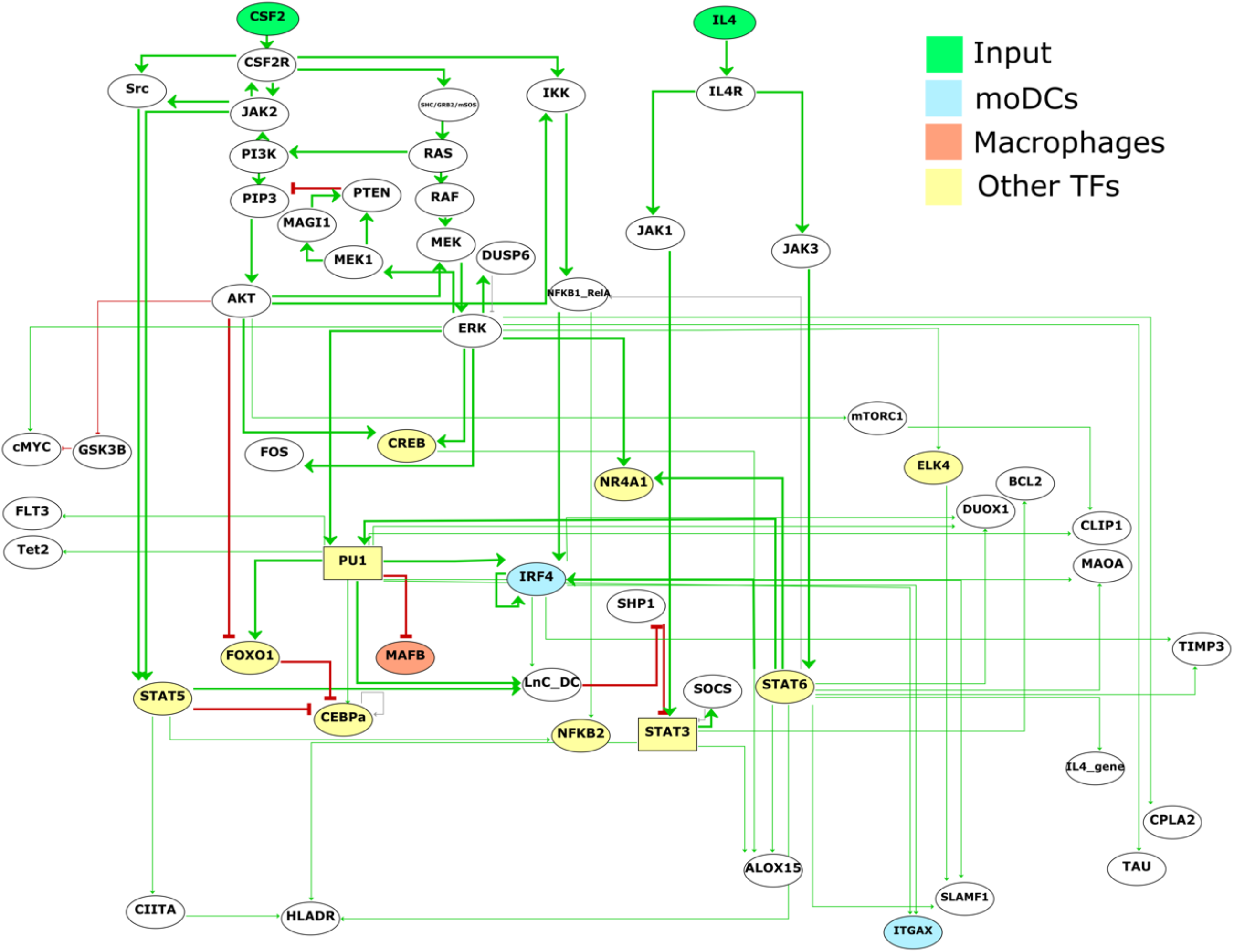
Regulatory graph controlling monocyte to moDCs differentiation, as derived from the scientific literature (last update “17april/2020”). The green nodes at the top represent the inputs (CSF2 and IL4), the yellow nodes denote transcription factors, the blue nodes denote moDCs specific genes, and orange nodes denote macrophage-specific genes. Nodes left in white correspond to components of generic signalling pathways. Green and red arcs denote positive and negative interactions, respectively.

Based on this first regulatory graph, we used GINsim to define logical rules (combining conditions on regulatory nodes with NOT, AND and OR Boolean operators), to compute the corresponding stable states and to perform simulations in order to determine the cellular phenotypes reached for each specific input condition. For this preliminary model, we found six stable states, but only one of them could be directly interpreted as a cellular phenotype (predenditic cells), while the other stable states did not recapitulate the typical signatures of activated monocytes or of macrophages.

Regarding the regulatory interactions between TFs and their target genes displayed in Figure 1, we observed that STAT6 has the highest number of interactions, while other TFs have only few interactions, such as STAT5, that only activates CIITA gene, or CREB that only activates ALOX15 gene. Furthermore, this regulatory graph contains very few specific moDCs markers. To complete this preliminary network, we decided to exploit epigenome and transcriptome data to infer novel regulatory interactions and integrate them into our logical model (a proof of concept of this approach can be found in Collombet *et. al.* 2016 (15)).

### Epigenome annotations help to unravel relevant transcription factor regulatory interactions

In order to complete our model of the regulatory network controlling the differentiation of monocytes into moDCs, we included the TFs known to be activated by CSF2 and IL4 signals in moDCs, as well as established monocytes markers. Moreover, we included information regarding the differentiation of monocytes into macrophages, which occurs when monocytes are treated with CSF2 alone (17). In short, we (i) used monocytes, moDCs, and macrophage epigenome data to define chromatin states, (ii) defined genomic regions likely to be involved in the regulation of the genes of the model, and (iii) searched for putative TFs binding sites in these regions.

We analysed ChIP-seq data from the Blueprint consortium for six histone marks (H3K4me1, H3K4me3, H3K27ac, H3K36me3, H3K9me3 and H3K27me3) in monocytes, moDCs, and macrophages derived from monocytes. We then used ChromHMM (18) to annotate the epigenome in each cell type based on these data. The resulting states (segments) were classified as Quiescent/low signal, Polycomb repressed, Poised regulation, Active TSS, Active promoter, Primed enhancer, Active gene/enhancer, Low transcription, TSS repressed and Strong transcription (Figure 2a). As expected, it is possible to visualize clear differences in the epigenome of moDCs and monocytes when exploring genes with specific cell expression in a genome browser, for example, the gene IRF4, a TF that mediates the differentiation of moDCs, is only active in moDCs while it is poised on macrophages and monocytes (Figure 2b).

**Figure 2.**
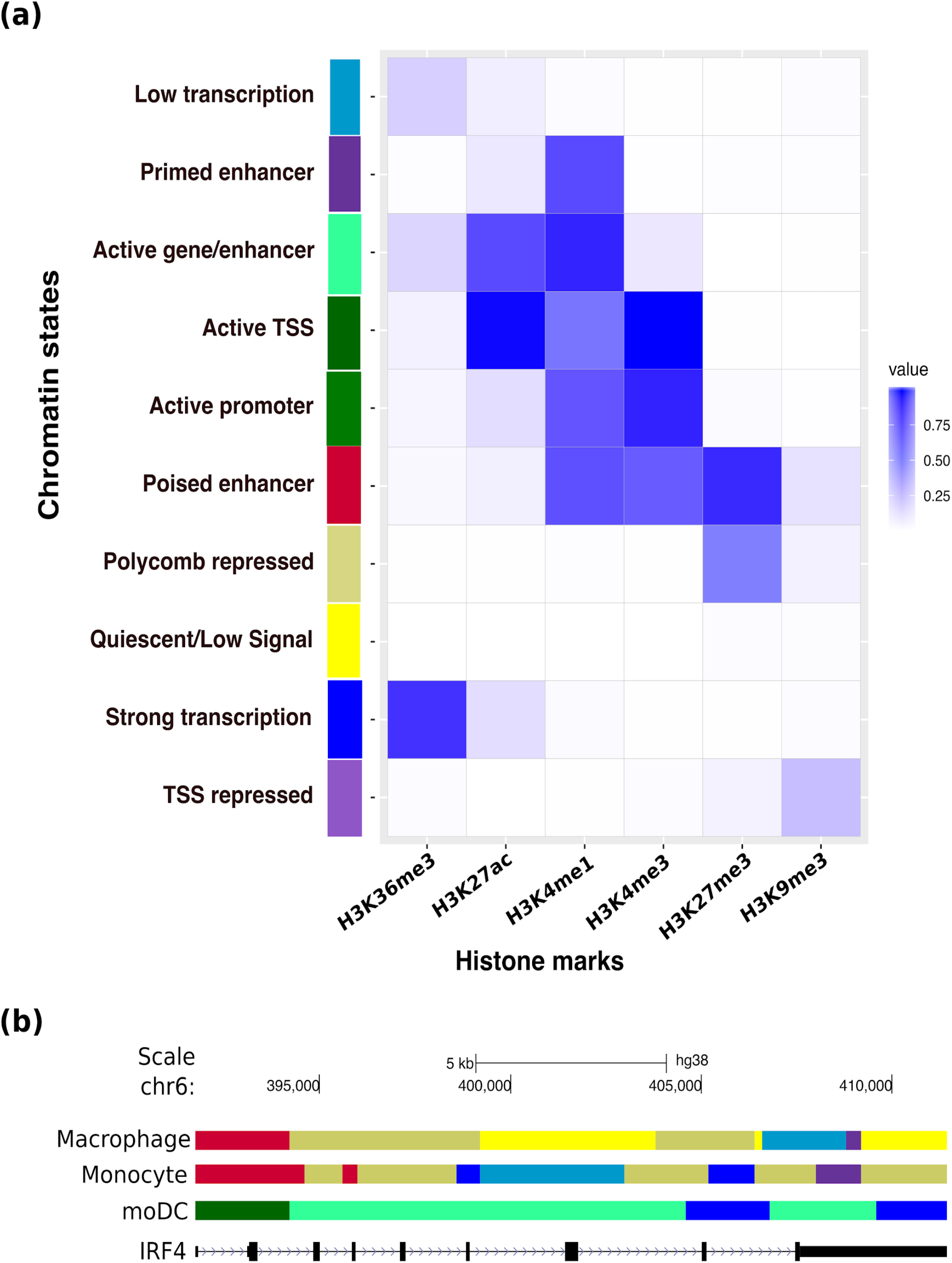
Epigenomic annotations of monocytes, moDCs, and macrophages. (a) Heatmap showing the histone mark enrichment in each of the states determined with ChromHMM. (b) IRF4 genomic region visualized in the UCSC browser, each row of segmentation corresponds to a specific cell type. Segmentation results from ChromHMM analysis, each color represents a state according to the color code used in the heatmap. State annotation was manually done based on biological knowledge. moDCs are the only cell type with the active gene marks (in green).

We selected the epigenome annotations regions with promoter-associated functions: Active TSS, Repressed TSS, Active gene/enhancer, and Poised regulation. These regulatory regions were then used to predict binding sites for the fourteen TFs activated by CSF2 and IL4 pathways (Figure 1) using the position-weight matrices collected in the Jaspar database (19) with the pattern-matching tool *matrix-scan* (20) from the RSAT suite (21). This led us to define novel regulatory interactions targeting specific gene markers for moDCs, monocytes, and macrophages (Table 1), thereby enabling us to complete the regulatory network controlling monocytes to moDCs differentiation.

**Table 1.** Cell type-specific gene markers selected to be added into the model. Based on the epigenome analysis we identified relevant regulatory interactions that helped select candidate genes to be added to the model.

| Cell-type | Gene |
| --- | --- |
| moDCs | TLR8 |
| moDCs | TLR7 |
| moDCs | TLR6 |
| moDCs | TLR4 |
| moDCs | TLR3 |
| moDCs | NCOR2 |
| moDCs | DEC205<br>(LY75) |
| moDCs | DCIR<br>(CLEC4A) |
| moDCs | CD83 |
| moDCs | CD48 |
| moDCs | CD226 |
| moDCs | CD209 |
| moDCs | CD1C |
| moDCs | CD1B |
| moDCs | CD1A |
| moDCs | CD141<br>(THBD) |
| moDCs | ITGAX<br>(CD11C) |
| moDCs | CCL22 |
| moDCs | CCL2 |
| Monocyte | CD14 |
| Monocyte | SELL |
| Macrophage | CD163 |
| Macrophage | CCDC151 |
| Macrophage | MERTK |
| Macrophage | CD206 |

For thirteen out of 20 genes related to moDCs phenotype, we found putative binding sites for the TF IRF4 (Figure 3a), corroborating a central role for IRF4 in the moDCs differentiation. In particular, we predicted that IRF4 directly regulates TLR genes (TLR3, 4 and 7), which play a crucial role in antigen recognition and are thus relevant for moDCs function. Furthermore, we predicted that TLR6 and TLR8 are regulated by STAT6, another essential TF in moDCs (6). In addition, we predicted that the genes encoding for the external proteins CD1A, CD1B, and CD1C are regulated by IRF4, as well as by other TFs (PU.1, PRDM1, NR4A1, CEBPA) related to moDC differentiation. Furthermore, we predicted that the gene coding for CD48, a costimulatory molecule involved in T cell activation, is regulated by PU.1, which is known to participate in the differentiation of STEM cell progenitors into leukocytes at different stages (22). We also looked for regulatory interactions between the identified TFs. Additionally, we validated interactions of PU.1 with CEBPA, IRF4, and IRF8. We also identified that AHR is regulating IRF4, MAFB, and PRDM1, which represent interesting candidates to assess experimentally. Figure 3b summarizes the regulatory interactions that compose our final logical model, where colored squares emphasize novel regulations, and asterisks denote interactions already described in the literature.

**Figure 3.**
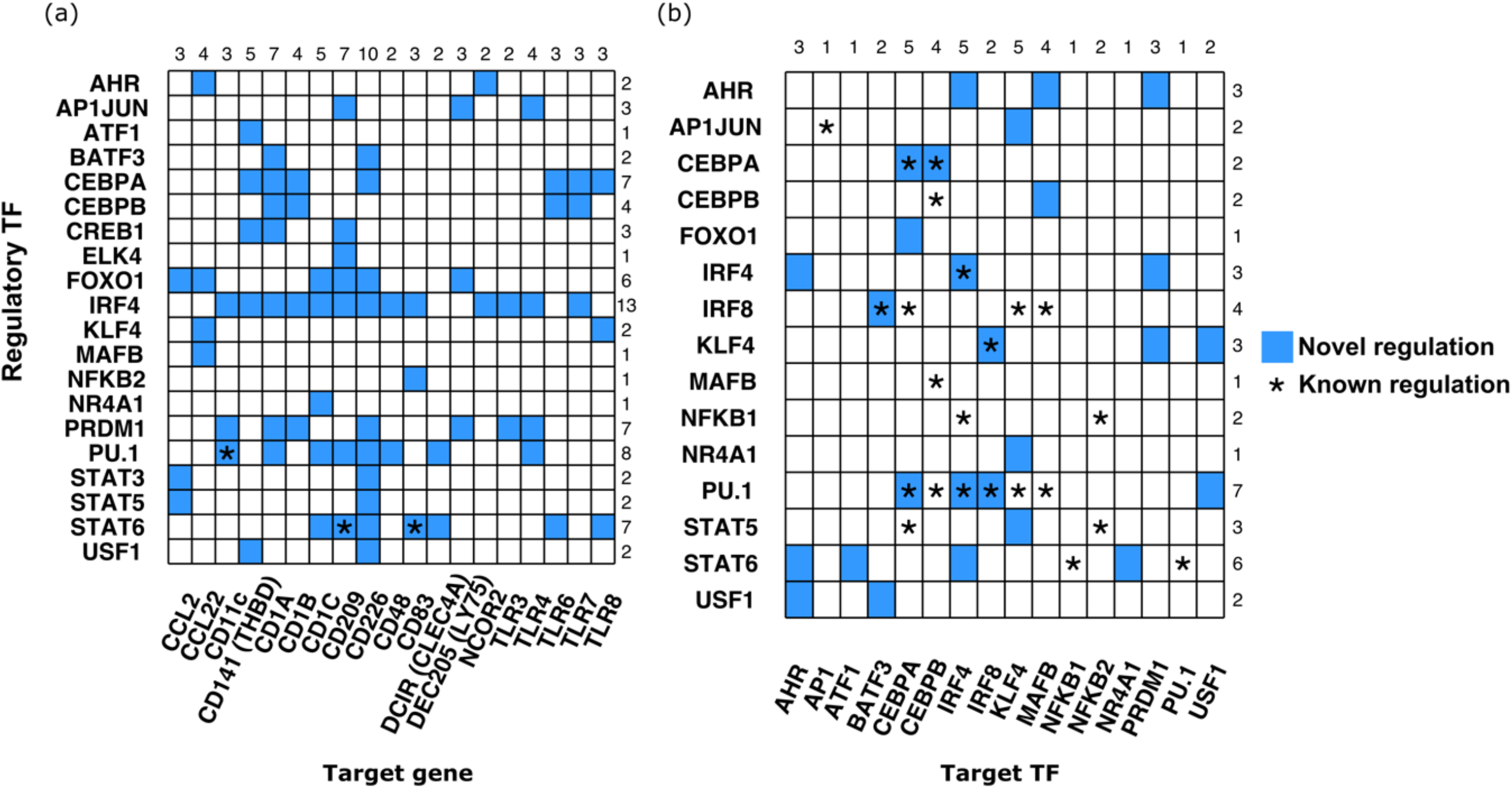
Predicted transcriptional regulatory interactions by TFs activated through CSF2 and IL4 signalling cascades. Each TF binding motif was used to search for putative binding sites in selected regulatory regions (based on chromatin state annotations) of specific gene or TF. (a) The y-axis shows the list of regulatory TFs and the x-axis shows the moDCs specific target genes. (b) The y-axis shows the list of regulatory TFs and the x-axis shows target TFs, in order to identify new regulatory interactions between TFs. Turquoise colored squares show predicted binding sites for the specified TF. Asterisks mark the binding sites for target genes or trans-regulation for target TF that have been already reported. Numbers at the end of each row correspond to the numbers of genes with regulatory interactions for each TF. Numbers at the top of every column correspond to the numbers of TF regulating each target gene.

### Integration of new relevant regulatory interactions improves model accuracy

We integrated the selected gene markers for each cell type with the predicted regulatory TFs into our model by adding the discovered regulatory interactions summarized in Figure 3. Using the new version of the model together with relevant Boolean rules, we set out to compute its stable states, which much better recapitulated the main cell fates compared to our first model (Figure 4a). Our revised model is characterised by four stable states. The first stable state corresponds to cell-death, which is the expected outcome for monocytes without cytokine stimulation. The second state, with IL4 ON, corresponds to monocyte signature (KLF4, SELL, and CD14 genes). The third stable state, with CSF2 ON, corresponds to monocytes that display a macrophage signature (MAFB, CEBPB, CD163, and CD206 genes). Finally, the last stable state, with CSF2 and IL4 ON, displays the moDCs signature (*i.e.* with IRF4, STAT6, CD1A, and CD209 all ON) (Figure 4b). Once we performed the analysis of stable states computation using our model, we validated that the PI3K signalling remained inhibited in order to reach moDCs commitment, this behavior was described by Van de Laar *et. al.* 2012 (23).

**Figure 4.**
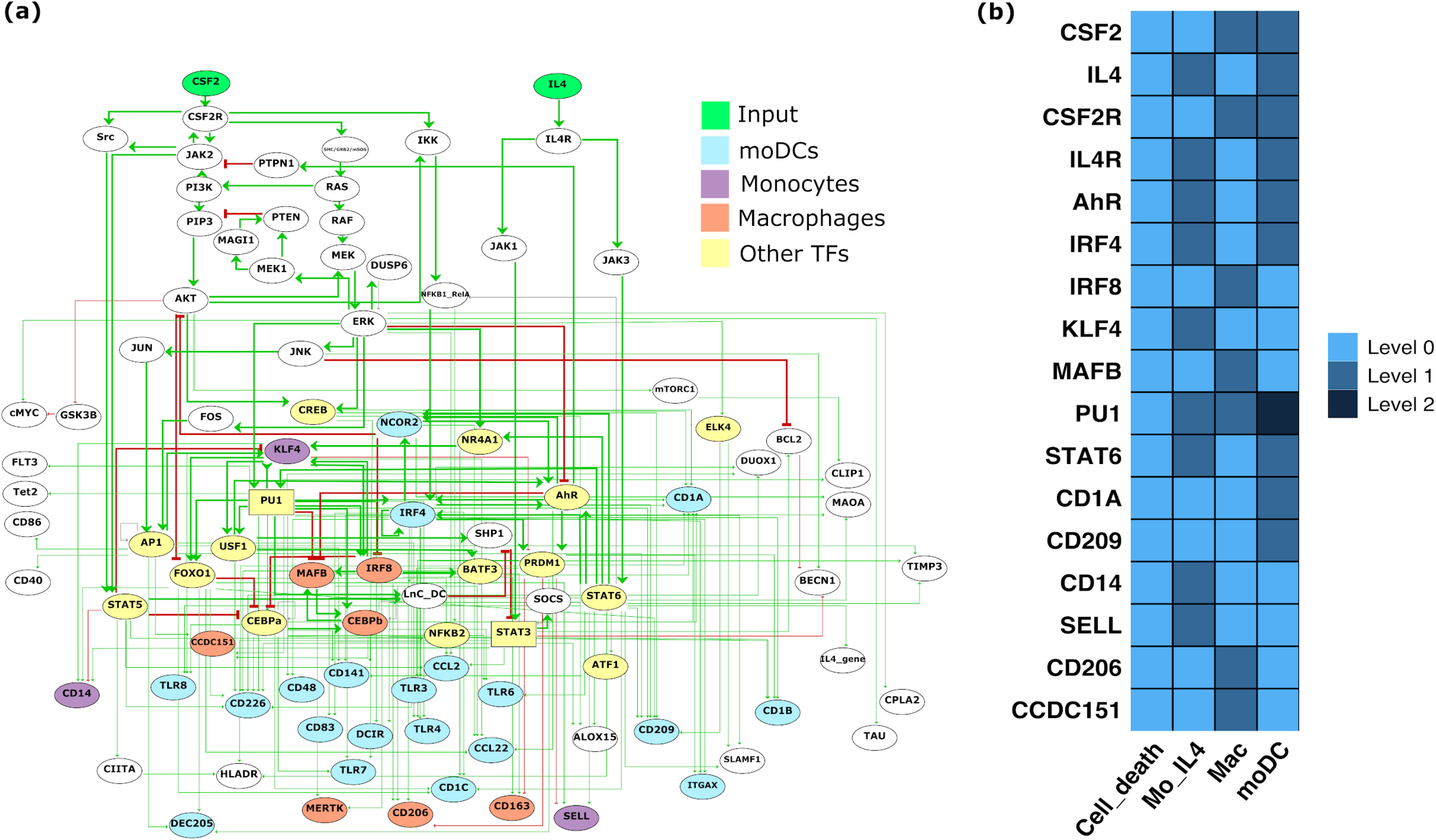
Logical model of monocytes to dendritic cells differentiation *in vitro*. **(a)** The green nodes at the top represent the inputs (CSF2 and IL4), the yellow nodes denote other TFs, blue nodes correspond to moDC specific genes, orange nodes to macrophage-specific genes, and purple nodes to monocyte specific genes. Green and red arcs denote positive and negative interactions, respectively. **(b)** Stables states of selected nodes (signature for each cell type), with the mention of the corresponding cell type. The first column corresponds to the final outcome in the absence of both IL4 and CSF2, *i.e.* cell-death of the monocytes. The second column corresponds to the stimulation of monocytes by IL4. The third column corresponds to the macrophage outcome, in the presence of the sole CSF2. Finally, the fourth column corresponds to moDCs commitment, in the presence of both IL4 and CSF2, where STAT3 reaches the level 2 in the presence of the long non-coding RNA LnC-DC, and PU.1 reaches the level 2, which is required to turn-off MAFB during moDCs commitment. SuppFig1 displays the complete set of nodes.

We used gene expression information to validate the different cell commitment expression signatures. To do that, we analysed RNA-seq data from monocytes, moDCs, and macrophages. Figure 5 displays the differential expression of the genes included in the model. Interestingly, we found two main clusters of genes highly expressed in moDCs, but down regulated in macrophages. These moDCs differentially express clusters of genes, including STAT3, STAT6, CEBPA, IRF4, TFs that participate in moDCs differentiation, also including CD206, MAOA, SLAMF1, that are specific markers for moDCs. Additionally, monocytes show highly expressed genes, like KLF4, IRF8, SELL, and CD14.

**Figure 5.**
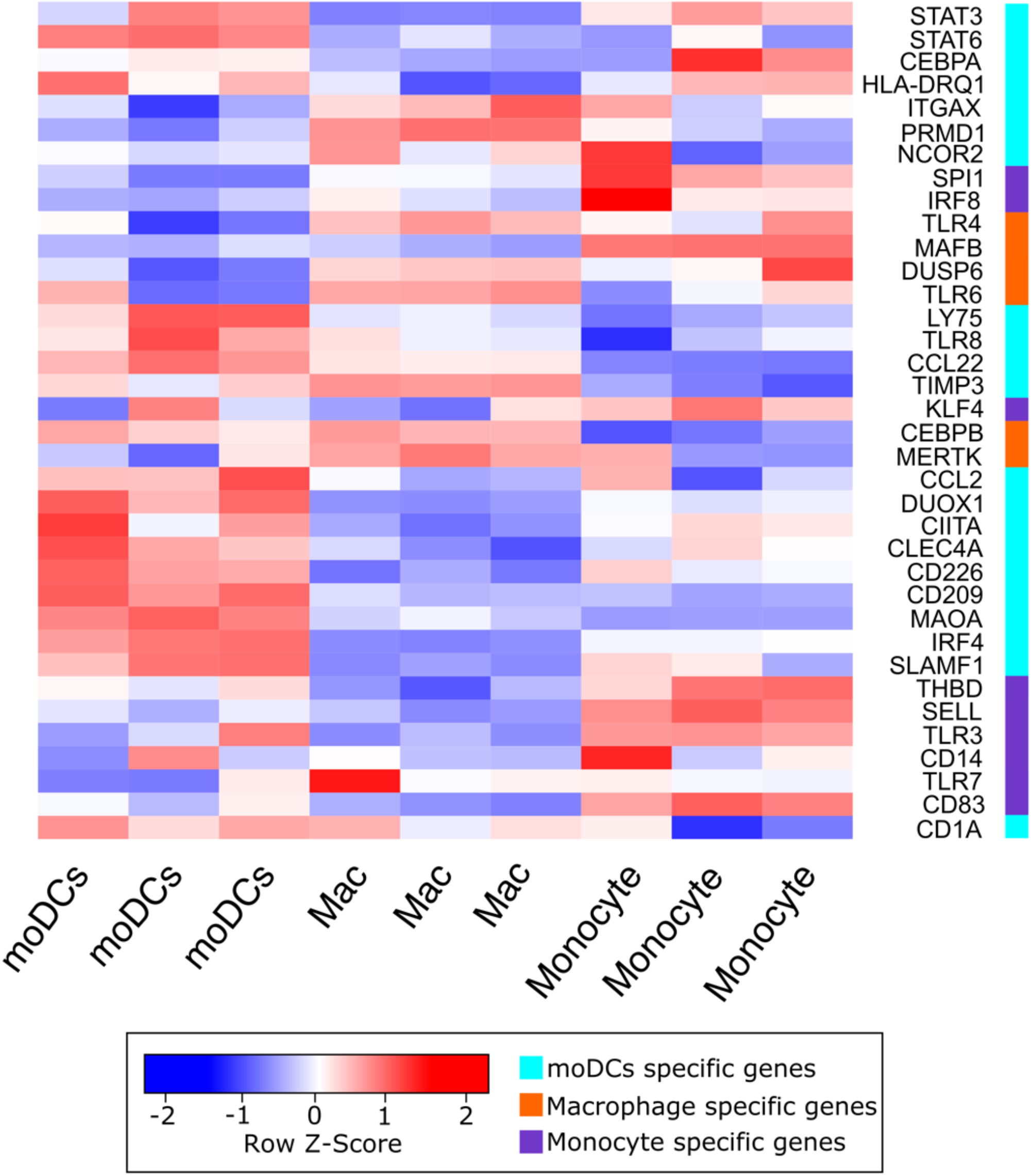
Clustered heatmap showing differentially expressed genes between cell types. The first three columns are moDCs, the next three columns are macrophages, and the last three columns are monocytes (columns represent biological replicates). The Z-score indicates the level of differential expression gene by gene bases. The colored column at the right represents specific gene markers per cell type, in blue for moDCs, orange for macrophages, and purple for monocytes. The heatmap is clustered by differential expression.

After integrating epigenome and transcriptome data into the model, we performed further simulations using the CoLoMoTo toolbox, with the purpose of recapitulating documented cellular commitment experiments.

### Model simulations correctly estimate cellular commitment to differentiation

We imported our model into the CoLoMoTo Interactive Notebook, a digital notebook that enables integrated complementary analyses software (with PINT, BioLQM, and MaBOSS) and facilitates reproducibility (24). The notebook is available as supplementary material. We used the tool Pint (25) to assess nine single gene mutants (IRF4, STAT6, PU.1, IRF8, MAFB, NCOR2, AHR, JAK3, CEBPB) that have been reported in the literature to affect the differentiation process. We were able to replicate the behavior of each mutant (perturbations) with our model. Table 2 shows the summary of the results obtained for these perturbations, while Figure 6 shows the behavior of each node for each perturbation at the corresponding stable states.

**Table 2.** Perturbations tested in the model of monocyte to moDCs differentiation.

| **Protein** | **Function** | **Phenotype described** | **Perturbation simulated** | **Model phenotype** |
| --- | --- | --- | --- | --- |
| **IRF4** | Transcription factor | Monocytes were infected using lentiviral vectors containing shRNA against IRF4, silenced IRF4 induced a dramatic reduction of moDCs (9). | Loss of function | Lack of most of moDCs specific markers |
| **STAT6** | Transcription factor | The ectopic expression of STAT6 in monocytes, resulted in increased levels of the DC-specific marker DC-sign, following CSF2 stimulation and without IL4 (6). | Gain of function | STAT6 is almost sufficient to archive moDC differentiation |
| **PU.1** | Transcription factor | Inducible constructions of PU.1 and MAFB were used to infected monocytes. In cells with PU.1 induced DCs, MafB differentiated macrophages (26). | Loss of function | Abolish moDCs and macrophage phenotype commitment |
| **IRF8** | Transcription factor | Introduction of KLF4 into an Irf8<sup>-/-</sup> myeloid progenitor cell line induced a subset of IRF8 target genes and caused partial monocyte differentiation (27). | Loss of function | Abolish KLF4 expression, and the entire macrophage differentiation |
| **MAFB** | Transcription factor | Silencing of MAFB resulted in a strong decrease in mo-Macs and an increase in mo-DC differentiation (26). | Loss of function | moDCs differentiation is normal, Macrophage differentiation is abolished |
| **NCOR2** | Transcriptional regulator | NCOR2 silencing resulted in 1,834 variable genes that correspond with IL4 signature genes (28). | Loss of function | Lack of moDCs specific markers |
| **AHR** | Transcription factor | AHR silencing reduced mo-DC differentiation while slightly increasing mo-Mac (9). | Loss of function | Lack of every moDCs specific markers, macrophage differentiation is normal |
| **JAK3** | Tyrosine-protein kinase | STAT6 phosphorylation disappeared following JAK3 inhibition. In the case of MACs, we did not observe STAT6 phosphorylation, given the lack of stimulation of JAK3 (6). | Loss of function | Macrophage phenotype with CSF2 and IL4. Macrophage differentiation is not affected |
| **CEBPB** | Transcription factor | In the absence of CEBPb in monocytes CEBPb-KO, only a very low amount (5%) of this macrophage-like morphology was seen, and most of the cells stayed round (29). | Loss of function | Lack of some specific macrophage markers |

**Figure 6.**
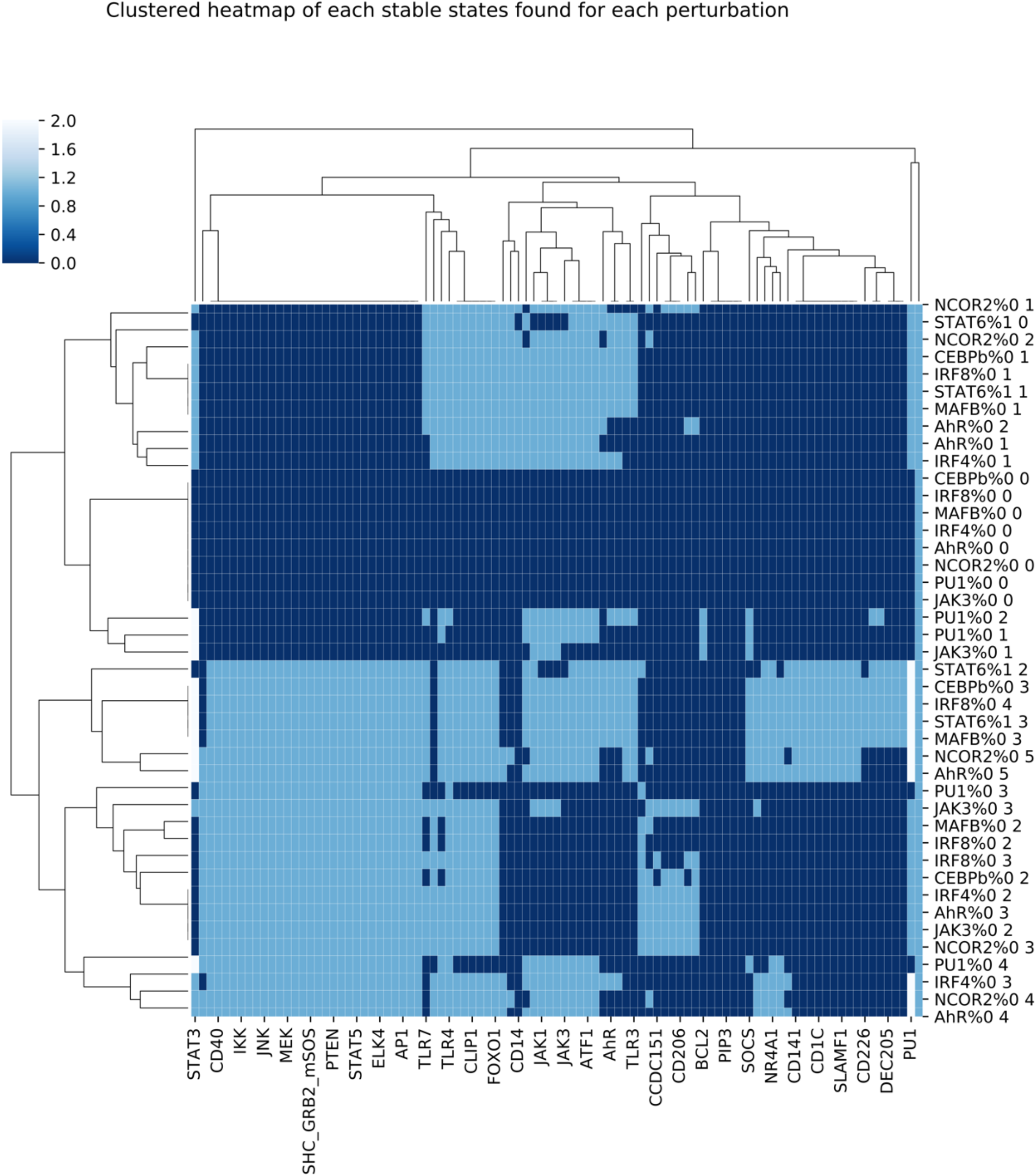
Clustered heatmap of the stable states obtained for all the perturbations considered. Each row represents one perturbation and one corresponding stable state. For example, NCOR2%0 1 denotes a knockout of NCOR2 and corresponds to the stable state 1 obtained for this condition. Every column represents one of the 95 nodes of the logical model.

We used the tool BioLQM to validate the reachability of each cell type commitment according to each stimulus combination. In order to verify the percentage of the final cell fate with the different initial stimulus, we used the stochastic Boolean simulation tool MaBoSS to estimate the probabilities to reach alternative states, where the final stable states represent alternative cell fate commitment (30).

From the literature, we know that CSF2 and ILF4 presence commits cells to differentiate to moDCs. Using MaBoSS, we tested cell commitment with the combination of CSF2 and IL4 ON at the initial state. This simulation showed that 100% of cells then commit to become moDCs (Figure 7c). For IL4 ON but not CSF2, 100% of the cells differentiate into stimulated monocytes, with the corresponding gene markers ON (figure 7a). For CSF2 ON but not IL4, 100% of the cells differentiated to macrophages, as expected (Figure 7b). In figure 7, we can clearly distinguish the three stable states corresponding to monocytes, moDCs, and macrophages, respectively.

**Figure 7.**
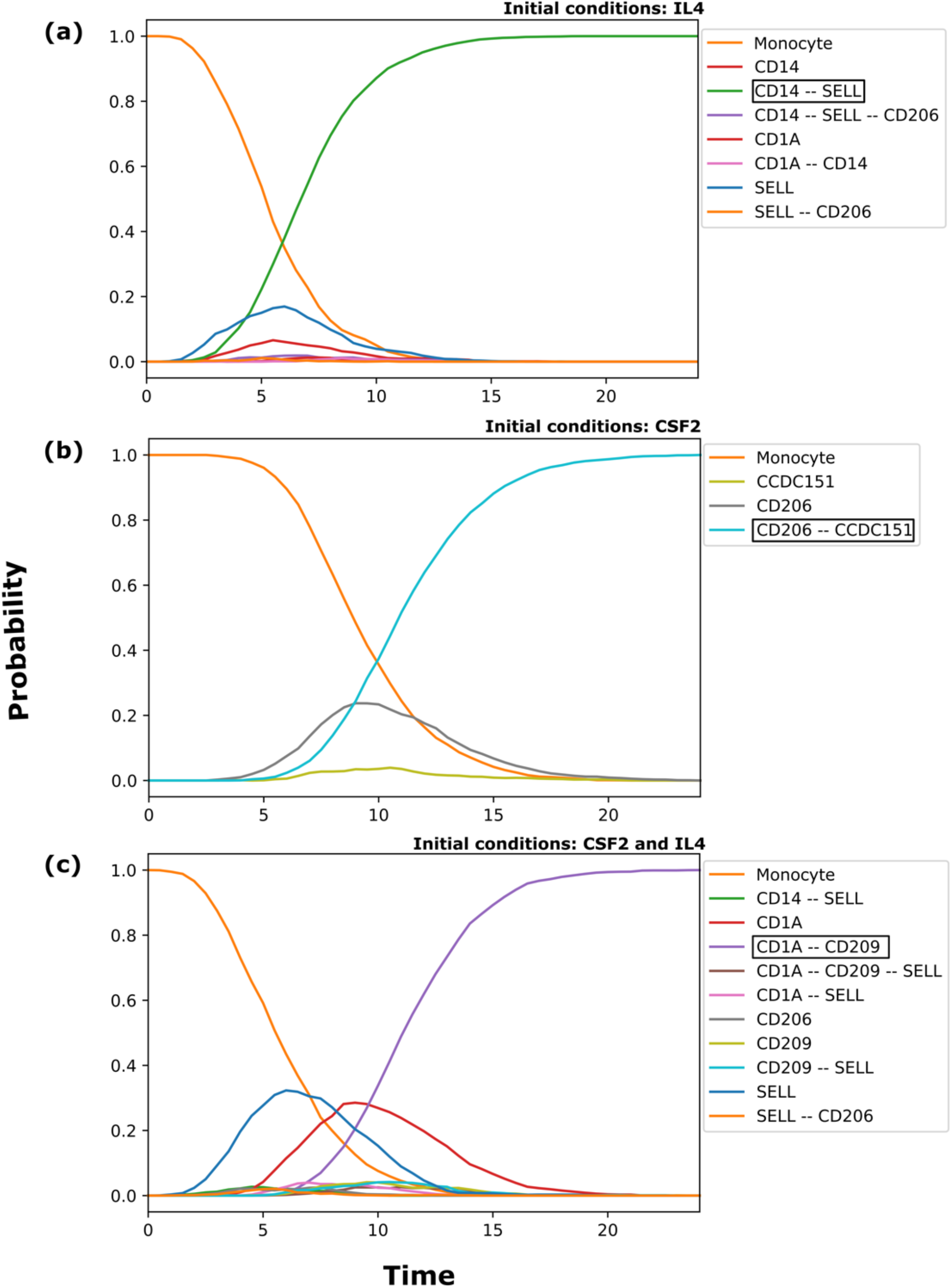
Stochastic simulations with. MaBoSS trajectories correctly recapitulate cellular commitment. The x-axis shows the time, the y-axis shows the probability to reach every final commitment. Highlighted in a rectangle is the final cellular commitment per stimulus. (a) IL4 ON gives rise to 100% differentiated cells into stimulated monocytes. (b) CSF2 ON gives rise to100% of the cells differentiated into macrophages. (c) With both CSF2 and IL4 ON 100% of cells commit to the moDC phenotype.

## Discussion

The construction of logical models traditionally relies on manual curation of the literature of a biological system of interest. In this work, we further took advantage of public ChIP-seq and RNA-seq data from the Blueprint consortium (31) to delineate in more detail the network driving the differentiation of monocytes into moDCs. We were able to fill various gaps in this regulatory network, which allowed us to reach a better understanding of this particular differentiation process. In the process, we predicted a series of novel interactions, validated *in silico* through our simulations, and amenable to further experimental tests.

In particular, we delineated a series of target genes presumably important for the differentiation of monocytes into moDCs. Some TFs are already well known, such as IRF4, AHR, STAT6, and PU.1 (6,9,32). In our analysis, we were able to recapitulate key features regarding the expression of the corresponding genes, such as a high expression of IRF4 and STAT6 genes in moDCs. We further validated the results obtained by Vento et al 2016 (6), in which STAT6 is required for moDCs differentiation; according to our model, STAT6 is indeed required for moDCs differentiation, but not for macrophage commitment.

We were also able to unravel TFs not previously reported as relevant in this process, such as FOXO1, C/EBPɑ, AP1, and PRDM1, tentatively regulating specific moDCs genes. We predict that FOXO1 regulates at least six moDCs genes, while C/EBPɑ regulates at least seven of them. Particularly, AP1 regulates TLR4, DEC205 (LY75), and CD209 (DC-SING), which are relevant for an antigen-presenting cell. CREB1 is also participating in the regulation of moDC genes, through the activation of CD141 y CD1A. We also predict for the first time that NR4A1 could regulate CD1C, a protein found at the surface of moDCs.

We further reviewed data recently published on the predicted TF-gene interaction considered in our model, and we found that some of these interactions have been recently experimentally confirmed. In particular, the regulation for the ITGAX gene was shown to be regulated by PU.1 and IRF4 (32), as predicted by our epigenomic analysis.

This study represents the first effort to integrate the current knowledge on monocytes to moDCs differentiation *in vitro* and should foster our understanding of this process. Additionally, we unraveled novel TFs and regulatory links potentially involved in this differentiation process.

## Material and methods

### GINsim implementation and simulations

Using the software GINsim version 3.0 (14), we integrated the previously described signalling pathways that are activated when monocytes are cultured with CSF2 and IL4 (studies reviews are inside GINsim model as annotations, and in the SuppFile1). We also performed a review of the available literature related to the process of monocyte to moDCs differentiation. The logical model was built using GINsim (33), where nodes represent genes or proteins, and edges represent the interactions between them, these interactions can be negative (red arrow) or positive (green arrow).

In general, each node can take two values, zero or one, but in special cases, activation (ON) requires to consider different qualitative levels of activation (*e.g.*: STAT3 expression is activated by JAK1, but the presence of LncDC leads to a further increase of STAT3 expression). For these special cases, it is possible to use multilevel nodes, *e.g.* ternary variables enabling an additional level of activation (hence taking the values 0, 1, and 2). Logical rules are associated with each component of the network, combining literals (*i.e.* regulatory variables with specific values) with the classical Boolean operators AND (&), OR (|) and NOT(!), thereby defining in which conditions each of these components can be activated or shut down.

### ChIP-seq data analysis

Raw fastq files from ChIP-seq experiments were retrieved from the Blueprint Consortium (31) data access portal (http://dcc.blueprint-epigenome.eu/#/datasets) with dataset identifiers EGAD00001001552, EGAD00001002484, EGAD00001002485, EGAD00001001576, EGAD00001002504. We processed six histone marks data (H3K4me1, H3K4me3, H3K27ac, H3K36me3, H3K9me3, and H3K27me3) with two biological replicates from human monocytes, macrophages, and moDCs. We performed quality control of read sequences with FastQC/0.11.3 tool (34), then we used Trimmomatic/0.33 (35) to improve quality reads before mapping them with bowtie2-2.2.6 (36) to the human hg38 reference genome. Second quality control is required after alignment, for which we used ENCODE QC, which consists of three major tests: NRF (non-redundant fraction), PBC1 (PCR Bottleneck coefficient 1), and PBC2 (PCR Bottleneck coefficient 2) (37). IDR analysis (37) was performed to replicate control.

### Chromatin states definition

We used one set of the six histone marks (H3K4me1, H3K4me3, H3K27ac, H3K36me3, H3K9me3, and H3K27me3) ChIP-seq data for each cell type (monocytes, macrophages, and moDCs) and their respective input control. Chromatin states were defined using ChromHMM (18) version 1.12 (38) with the recommended parameters (BinarizeBed -b 200, assembly hg38), and specifying 10 states. In order to define the description for the states, we used the probability of appearance of different marks in every state (*e.g.* H3K27ac-Enhancers, H3Kme1-Enhancers, H3K4me3-Promoters, H3K27me3-Repressive, H3K9me3-Repressive, H3K36me3-Transcribed (39), and then we looked into the enrichment of the states for several genome annotations (CpGIsland, RefSeqExon, RefSeqGene, RefSeqTES, RefSeqTSS, and RefSeq2kb). Integrating this information, we were able to manually assign a functional description to each state. Once we described every state, we focused on Active TSS, Repressed TSS, Active gene/enhancer, and Poised regulation regions to look for regulatory interactions between TFs and target genes. For poised regulation, it is well known that regions go from poised to active regions when cells are under differentiation. Additionally, we took the whole segment for each state selected result from ChromHMM.

### Search for TFBS using matrix-scan

From manual curation of literature, we identified 22 TFs that participate after monocyte stimulation leading to the differentiation of macrophages or moDCs. We retrieved one PSSM (Position-Specific Scoring Matrix) for each of the 22 TFs (SuppTable1) from the JASPAR2018 database human collection (19). We performed pattern-matching searches for TF motif instances using the 22 PSSMs in the selected chromatin regions (Active TSS, Repressed TSS, Active gene/enhancer, and Poised regulation) from ChromHMM results. For this task we used the tool *matrix-scan* (20) from the RSAT suite (21) with the following main parameters: background model of Markov order 1 and stringent thresholds of p-value ≤ 10^−5^ and score 1 (-markov 1 -lth score 1 -uth pval 1e-5).

### RNA-seq analyses

Raw fastq files from RNA-seq experiments were retrieved from the Blueprint Consortium (31) data access portal (http://dcc.blueprint-epigenome.eu/#/datasets) with dataset identifiers: EGAD00001002308, EGAD00001001506, EGAD00001002526, EGAD00001002507, and EGAD00001001582. For this analysis, we used the methods described in Law et al 2016 (40). In brief, that is quality control with FastQC/0.11.3 (41), pseudo-alignment and count determination with Kallisto 0.43.1 (42) using the release-90 from Ensembl (ftp://ftp.ensembl.org/pub/release-90/fasta/homo_sapiens/cdna/Homo_sapiens.GRCh38.cdna.all.fa.gz) to create our index with the following command: kallisto index -i index_kallisto_hsap_90_cdna --make-unique Homo_sapiens.GRCh38.cdna.all.fa.gz. Counts were assigned to genes using Tximport 1.14.0 (43), and were processed from raw-scale to counts per million (CPM), then they were transformed to log-CPM. Genes below 1 of expression were removed. Then we normalized raw library sizes using the *calcNormFactors* function from edgeR library in R. Afterwards, we performed a differential gene expression analysis with edgeR 3.28.0 (44). Finally, we used heatmap.2 from the gplots library to plot the genes found in our model (Figure 5).

### CoLoMoTo analysis

In order to assure reproducibility, we used the CoLoMoTo toolbox(16) that integrates several logical modeling tools, including GINsim, bioLQM, Pint, and MaBoSS. We used GINsim to compute the stable states, and bioLQM to identify trap spaces approximating cyclic attractors. The computation of mean stochastic trajectories was performed using MaBoSS (30). The GINsim model and the CoLoMoTo notebook are available at https://github.com/karenunez/moDC_model_differentiation.

### Figures generation

Figure 1, and 4A were generated with the GINsim software. The plots in Figures 2A, 3A, 3B, and 4B were done using the ggplot2 library from R. Figures 6, and 7 are from the CoLoMoTo notebook constructed in this study.

## Supporting information

SuppFile Model annotation

SuppMaterial

## Supplemental material

Supplementary files are available at https://github.com/karenunez/moDC_model_differentiation.

SuppFile1. Model_annotation.doc

SuppFile2. Mo_Mac_moDCs_ChromHMM_summary.html

SuppFile3. moDC_E7_ActiveGeneEnhancer.bed

SuppTable1. TFs_JASPARID_matrixes.xlsx

SuppFigure1.StableStates_95nodes.png

## Acknowledgments

Karen Julia Nuñez Reza is a doctoral student from Programa de Doctorado en Ciencias Biomédicas, Universidad Nacional Autónoma de México (UNAM) and received fellowship CVU/Becario: 634764/331823 from CONACYT, México. This work received support from Luis Aguilar, Alejandro Leon and Jair García of the Laboratorio Nacional de Visualización Científica Avanzada. We thank Carina Uribe Díaz and Alejandra Castillo Carbajal for their technical support. We want to thank Marc Dalod and Thien Phong Vu Manh for your valuable comments on our work.

## Authors’ contributions

K.J.N.-R. carried out the manual literature curation, performed the ChIP-seq and RNA-seq data analysis, participated in the construction of the two model versions, notebook implementation, study design, and drafted the manuscript; A.N. participated in the construction of the two model versions, notebook implementation, and drafted the manuscript; A.S.-J. participated in the manual literature curation, and drafted the manuscript; A.V.L.-A. participated in the manual literature curation, and drafted the manuscript; A.S. participated in the design of the study, and drafted the manuscript; M.T.C. mentored the ChIP-seq, ChromHMM, and matrix-scan analysis, participated in the design of the study and drafted the manuscript; D.T participated in the construction of the two model versions, notebook implementation, study design, and drafted the manuscript; A.M.-R. mentored the ChIP-seq, ChromHMM, and matrix-scan analysis, participated in the design of the study, and drafted the manuscript.

