## Supplementary material for "Logical modeling of dendritic cells *in vitro* differentiation from human monocytes unravels novel transcriptional regulatory interactions": SuppFile Model annotation

Description of the model "Karen_MoDC_16april2020"

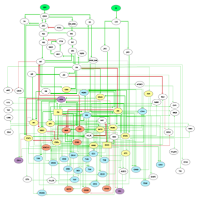

Annotation

Sallusto and Lazavecchia in 1994 describe for the first time the in vitro protocol for human monocyte to dendritic cell (moDC) differentiation using granulocyte macrophage colony stimulating factor (CSF2) and interleukin 4 (IL4).

In this work we integrated scientific report knowledge of the signaling cascades that initiated the differentiation process, we also integrated new possible transcriptional regulation result from the analysis of regulatory regions from monocyte, dendritic cells and macrophages as negative control of monocyte to moDC, macrophages results after the stimulation of monocytes with CSF2.
https://www.ncbi.nlm.nih.gov/pubmed/8145033

Nodes

| ID | Logical function |
|---|---|
| CSF2<br>Sallusto and Lazavecchia in 1994 describe for the first time the in vitro protocol for human monocyte to dendritic cell (moDC) differentiation using granulocyte macrophage colony stimulating factor (CSF2) and interleukin 4 (IL4).<br>1. https://www.ncbi.nlm.nih.gov/pubmed/8145033 | Input node |
| IL4<br>Sallusto and Lazavecchia in 1994 describe for the first time the in vitro protocol for human monocyte to dendritic cell (moDC) differentiation using granulocyte macrophage colony stimulating factor (CSF2) and interleukin 4 (IL4).<br>1. https://www.ncbi.nlm.nih.gov/pubmed/8145033 | Input node |
| CSF2R<br>PMID:22323450. The GM-CSFR contains 2 distinct subunits, the GM CSF specific-chain (GM CSFR; CD116) and the common receptor, which is shared between the GM CSFR, the IL3 receptor,and the IL5 receptor. Downstream signaling cascades areprimarily induced through interaction of effector proteins with the BetaC subunit. Signaling is initiated by the cytoplasmic tyrosine kinasejanus kinase 2 (JAK2), which then acts on various downstreamproteins.<br>1. https://www.ncbi.nlm.nih.gov/pubmed/22323450 | • CSF2 |
| IL4R<br>PMID: 25159217. The IL4 receptor (IL4R) signals the activation of the Janus kinase 3 (JAK3) STAT6 pathway through its common gamma chain, which leads to the development of immature DCs.<br>1. https://www.ncbi.nlm.nih.gov/pubmed/25159217 | • IL4 |

| | |
|---|---|
| AHR<br><br>PMID: 23430108. Whereas ERK inhibition upregulated AhR dependent transcriptionin in vitro generated monocyte derived cells (moDCs and Macrophages).<br><br>FOXO and AhR autoregulation came from putative TFBS perform with matrix-scan with all the matrixes of every TF in the model.<br>1. https://www.ncbi.nlm.nih.gov/pubmed/23430108 | • (NCOR2 & USF1 & STAT6) \| (IRF4 & STAT6 & !ERK) |
| AP1<br>Macrophage specific H3K4me1 regions were characterized by a distinct motif composition, including GT box, an AP1 like motif, an E box element, the consensus PU.1 motif, a composite CEBP_bZIP element, and a NFKB motif.<br>1. https://pubmed.ncbi.nlm.nih.gov/22550342 | • JUN & FOS |
| ATF1<br>STAT6<br>is a putative regulation, through the sites found using matrix-scan with the regulatory regions of moDC in this study. | • STAT6 |
| BATF3<br>PMID: 28781277. KLF4 and BATF3 serve as critical transcription factors downstream of IRF8 to induce the differentiation of monocytes and DCs, respectively.<br><br>USF1 is a putative regulation, through the sites found using matrix-scan with the regulatory regions of moDC in this study.<br>1. https://www.ncbi.nlm.nih.gov/pubmed/28781277 | • USF1 \| IRF8 |
| CEBPa<br><br>PMID: 28781277. Conversely, IRF8 blocks the activity of the transcription factor CEBPalfa to suppress the neutrophil differentiation program.<br><br>CEBP transcription factors, and CEBPb in particular, have long been implicated in the regulation of monocyte macrophage differentiation, whereas CEBPa appears to be more important for the maturation of granulocytes<br>1. https://www.ncbi.nlm.nih.gov/pubmed/28781277<br>https://pubmed.ncbi.nlm.nih.gov/21558273/ | • PU1 & !FOXO1 & !IRF8 & !STAT5 |
| CEBPb<br>Macrophage specific H3K4me1 regions were characterized by a distinct motif composition, including GT box, an AP1 like motif, an E box element, the consensus PU.1 motif, a composite CEBP_bZIP | • PU1 & (CEBPa \| MAFB) |

element, and a NFKB motif.

CEBP transcription factors, and CEBPb in particular, have long been implicated in the regulation of monocyte macrophage differentiation, whereas CEBPa appears to be more important for the maturation of granulocytes

1. https://pubmed.ncbi.nlm.nih.gov/22550342

https://pubmed.ncbi.nlm.nih.gov/21558273/

| | |
|---|---|
| FOS<br>PMID: 23430108. U0126 treatment significantly reduced the expression of wellknown targets of ERK (FOS, MYC, DUSP6).<br>1. https://www.ncbi.nlm.nih.gov/pubmed/23430108 | • ERK |
| cMYC<br>PMID:22323450.<br>PMID: 23430108. U0126 treatment significantly reduced the expression of wellknown targets of ERK (FOS, MYC, DUSP6).<br>1. https://www.ncbi.nlm.nih.gov/pubmed/22323450<br>https://www.ncbi.nlm.nih.gov/pubmed/23430108 | • ERK & !GSK3B |
| CREB<br>MAPK signaling pathway Homo sapiens<br>1. https://www.genome.jp/dbget-bin/www_bget?hsa04010 | • AKT \| ERK |
| ELK4<br>MAPK signaling pathway Homo sapiens<br>1. https://www.genome.jp/dbget-bin/www_bget?hsa04010 | • ERK |
| FOXO1<br>PMID:22323450. Activated PKB regulates many targets, including the FOXO transcription factors, the TSC1 TSC2 complex, and the mTOR complex 1 (mTORC1).<br><br>KLF4 is a putative regulation, through the sites found using matrix-scan with the regulatory regions of moDC in this study.<br>1. https://www.ncbi.nlm.nih.gov/pubmed/22323450 | • (PU1 \| KLF4) & !AKT |
| IRF4<br><br>DCs were found to express IRF4 mRNA and protein constitutively, and STAT and NFkB transcription factors play an important role inregulating IRF4 expression in DCs. IRF4 protein bound to a regulatory element in its own promoter, suggesting an autoregulatory loop controlling IRF4 mRNA expression in DCs.<br><br>AhR is a putative regulation, through the sites found using matrix-scan with the regulatory regions of moDC in this study.<br><br>Monocytes stimulated with IL4 activates IRF4. | • AhR \| (PU1 & STAT6 & NFKB1_RelA & IRF4) |

1. https://pubmed.ncbi.nlm.nih.gov/10453013
https://pubmed.ncbi.nlm.nih.gov/29871928/

IRF8

• (PU1 | KLF4) & !NCOR2

PMID: 29262348. We found a statistically significant enrichment of genes upregulated in the IL4 signature in MOs GMCSF IL4(0-72h) treated with scrambled siRNAs (Figure6I) whereas genes downregulated in the IL4 signature were enriched in MOs GMCSF IL4(0-72h) treated with antiNCOR2 siRNAs (Figure 6J), establishing NCOR2 as a key regulator for IL4 induced MO differentiation.

NCOR2 is a putative negative regulation, through the sites found using matrix-scan with the regulatory regions of moDC in this study.

1. https://www.ncbi.nlm.nih.gov/pubmed/29262348

KLF4

• (NR4A1 & IRF8 & AP1) | (PU1:1 & !STAT5)

PMID: 17762869. PU.1 induced the KLF4 promoter 15 fold.

1. https://www.ncbi.nlm.nih.gov/pubmed/17762869

MAFB

• (CEBPb | IRF8) & !AhR & !PU1:2

PMID: 24070385. Pu.1 in monocytes favors DC development at the expense of a macrophage fate by directly inhibiting expression of the macrophage factor, MafB, suggesting that Pu.1 could be an important decision factor between DC and macrophage commitment.

PMID: 22868453. MafB gene silencing improved the differentiation potential of CD14+ cells into mDCs, increasing the percentage of mDCs by >75%. Furthermore, GATA-1+ and HLA-DR+ mDCs were increased following MafB silencing.

We have shown that the transcription factors MafB and PU.1 induce alternative macrophage or DC fates respectively in myeloblasts.

1. https://www.ncbi.nlm.nih.gov/pubmed/24070385
https://www.ncbi.nlm.nih.gov/pubmed/22868453
https://pubmed.ncbi.nlm.nih.gov/28338898/

NFKB1_RelA

• IKK

PMID:22323450. Activation is achieved through the IKK complex, which phosphorylates IkB proteins. These are subsequently ubiquitinated and finally degraded, enablingnuclear translocation of canonical NFkB dimers.

1. https://www.ncbi.nlm.nih.gov/pubmed/22323450

NFKB2

• NFKB1_RelA | STAT5

PMID:22323450. In the previous section, GM CSF induced activation of STAT5 and canonical NFkB transcription factors increases the intrinsic immunogenicity of the DCs generated

1. https://www.ncbi.nlm.nih.gov/pubmed/22323450

NR4A1

- ERK & STAT6

MAPK signaling pathway Homo sapiens

1. https://www.genome.jp/dbget-bin/www_bget?hsa04010

PRDM1

- IRF4 & AhR & !KLF4

PMID: 28930664. PRDM1 silencingsignificantly decreased moDC differentiation, while increasingthe proportion of moMacs (Figure 5D).

IRF4 & KLF4 regulation is due a putative TFBS found with matrix-scan

1. https://www.ncbi.nlm.nih.gov/pubmed/28930664

PU1

- (ERK | STAT6) & !(ERK & STAT6)

2

PMID: 20510871. One study concluded that PU.1 is necessary for all DC development because Sfpi1 homozygotic mutant neonates lacked thymic DCs and were unable to generate DCs in vitro in response to GMCSF.

We have shown that the transcription factors MafB and PU.1 induce alternative macrophage or DC fates respectively in myeloblasts.

IL4 induced threonine phosphorylation of PU.1, which was susceptible to these inhibitors. Because it is known that phosphorylation of PU.1 at serine 148, located within a casein kinase II consensus motif.

1. https://www.ncbi.nlm.nih.gov/pubmed/20510871
https://pubmed.ncbi.nlm.nih.gov/28338898/
https://pubmed.ncbi.nlm.nih.gov/11594748/

STAT3

- JAK1 & SHP1

2

However, by far the most important negative regulation occurs at the level of receptor mediated STAT3 activation and is conferred by suppressor of cytokine signalling (SOCS) E3 ubiquitin ligases that enable the degradation of cytokine receptor complexes. SOCS proteins are themselves encoded by STAT target genes and thus provide a transcription dependent negative feedback mechanism.

lncDC bound directly to STAT3 in the cytoplasm, which promoted STAT3 phosphorylation on tyrosine705 by preventing STAT3 binding to and dephosphorylation by SHP1.

1. https://www.ncbi.nlm.nih.gov/pubmed/30578415
https://www.ncbi.nlm.nih.gov/pubmed/27165851
https://www.ncbi.nlm.nih.gov/pubmed/28465674
https://www.ncbi.nlm.nih.gov/pubmed/24744378

STAT5 :• Src | JAK2

PMID:22323450. Of the signaling proteins contributing to GM CSF driven DC development, JAK2 activated STAT5 is the clearest regulator of differentiation.

1. https://www.ncbi.nlm.nih.gov/pubmed/22323450

STAT6 :• JAK3

The IL4 receptor (IL4R) signals the activation of the Janus kinase 3 (JAK3) STAT6 pathway through its common gamma chain, which leads to the development of immature DCs.

1. https://www.ncbi.nlm.nih.gov/pubmed/25159217
https://www.ncbi.nlm.nih.gov/pubmed/10485906
https://www.ncbi.nlm.nih.gov/pubmed/16540365

USF1 :• PU1 | KLF4

Involvement of USF in myeloid cell differentiation was suggested by Kreider et al

KLF4 is a putative regulation, through the sites found using matrix-scan with the regulatory regions of moDC in this study.

1. https://pubmed.ncbi.nlm.nih.gov/10085160

NCOR2 :• IRF4 | AhR | STAT6

AhR and IRF4 regulation is result from the matrix-scan analysis

We identified NCOR2 as a key transcriptional hub linked to IL4 dependent differentiation of MOs.

1. https://pubmed.ncbi.nlm.nih.gov/29262348

JAK2 :• CSF2R & !PTPN1

PMID:22323450. Src kinases are recruited to BetaC by their SH2 domains that interact with phosphorylated Y612, Y695, and Y750. The STATs are primarily phosphorylated by JAK2, but kinase activity of the Src kinases has also been reported.

PMID: 11694501. In this study, we have shown that PTP1B recognizes TYK2 and JAK2, but not JAK1, and can modulate signaling responses to IFNgamma and IFNalfa

1. https://www.ncbi.nlm.nih.gov/pubmed/22323450

Src :• CSF2R:1 | JAK2

PMID:22323450. Src kinases are recruited to BetaC by their SH2 domains that interact with phosphorylated Y612, Y695, and Y750. The STATs are primarily phosphorylated by JAK2, but kinase activity of the Src kinases has also been reported.

1. https://www.ncbi.nlm.nih.gov/pubmed/22323450

PI3K :• RAS | JAK2

PMID:22323450. Activity of PI3K is promoted by JAK2 mediated phosphorylation of p85

1. https://www.ncbi.nlm.nih.gov/pubmed/22323450

PIP3 :• PI3K & !PTEN

PI3K functions mainly through the generation of PIP3, an activity counteracted by phosphatases PTEN and SHIP.

1. https://www.ncbi.nlm.nih.gov/pubmed/22323450

AKT :• PIP3 & !NCOR2

PMID:22323450. PIP3 acts as a second messenger, regulating a large variety of downstream targets, including protein kinase B (PKB; also called AKT).

Promotes activation of phosphatidylinositol 3 kinase and of the AKT1 signaling cascade

1. https://www.ncbi.nlm.nih.gov/pubmed/22323450
https://www.genecards.org/cgi-bin/carddisp.pl?gene=PTK2B&keywords=PTK2B

PTEN :• MAGI1 & MEK1

MAGI1 contains six PDZ domains, a guanylate kinase, GUK, domain, and two WW domains flanked by two PDZ domains. PTEN binds selectively to the second PDZ domain of MAGI1. PTEN interacted indirectly with beta_catenin by binding the scaffolding protein MAGI1b. scaffolding molecules such as MAGI1b, which was reported to interact with beta catenin

We report the existence of a ternary complex between MEK1, MAGI1, and PTEN, mediating the translocation of PTEN to the membrane and

therefore regulating the concentration of PIP3 and AKT activation.
Both MEK1 and MAGI1 are necessary for complex formation, and PTEN will not bind to one component if the other is missing.
1. https://www.ncbi.nlm.nih.gov/pubmed/22323450
https://www.ncbi.nlm.nih.gov/pubmed/23453810

MEK1 : • ERK
PMID: 23453810. Thus, phosphorylation of MEK1 T292 relays a negative feedback within the ERK pathwayand initiates the deactivation of the PIP3 AKT pathway through the membrane localization of MAGI PTEN, acting as a temporal switch for both cascades. Thus, ERK differentially
regulates binding of MEK1 to WW domain containing proteins and may negatively affect survival by promoting the translocation of WOX1 to the mitochondria (Lin et al., 2011) and the membrane recruitment of PTEN in the context of the MEK1/ MAGI1/PTEN complex.
1. https://www.ncbi.nlm.nih.gov/pubmed/23453810

MAGI1 : • MEK1
PMID: 23453810. Thus, MEK1 is essential for the formation of a complex containing MAGI1 and PTEN and for their membrane translocation upon growth factor stimulation. Mutation of the WW domains of MAGI1, in particular of WW2, strongly reduced MEK1 binding by a WW MAGI1 fragment or by full length MAGI1.
We report the existence of a ternary complex between MEK1, MAGI1, and PTEN, mediating the translocation of PTEN to the membrane and therefore regulating the concentration of PIP3 and AKT activation.
Both MEK1 and MAGI1 are necessary for complex formation, and PTEN will not bind to one component if the other is missing.
MEK1 ablation prevented MAGI1 membrane translocation
1. https://www.ncbi.nlm.nih.gov/pubmed/23453810

CLIP1 : • mTORC1 | PU1
mTOR signaling pathway Homo sapiens
1. https://www.genome.jp/dbget-bin/www_bget?hsa04150

mTORC1 : • AKT
PMID: 22323450. Activated PKB regulates many targets, including the FOXO transcription factors, the TSC1 TSC2 complex, and the mTOR complex 1 (mTORC1).
PMID: 27614799
1. https://www.ncbi.nlm.nih.gov/pubmed/22323450

SHC_GRB2_mSOS : • CSF2R
PMID:22323450. The principle MAPK pathway activated by the GM CSF receptor is the MEK ERK pathway. Recruitment of mSOS to the SHC GRB2 complex enables mSOS to catalyze RAS activation.

1. https://www.ncbi.nlm.nih.gov/pubmed/22323450

RAS • SHC_GRB2_mSOS

PMID:22323450. The principle MAPK pathway activated by the GM CSF receptor is the MEK ERK pathway. Recruitment of mSOS to the SHC GRB2 complex enables mSOS to catalyze RAS activation. Formation of active GTP bound RAS from inactive GDP bound RAS leads to the successive activation of RAF, MEK, and ERK.

1. https://www.ncbi.nlm.nih.gov/pubmed/22323450

RAF • RAS

PMID:22323450. The principle MAPK pathway activated by the GM CSF receptor is the MEK ERK pathway. Recruitment of mSOS to the SHC GRB2 complex enables mSOS to catalyze RAS activation. Formation of active GTP bound RAS from inactive GDP bound RAS leads to the successive activation of RAF, MEK, and ERK.

1. https://www.ncbi.nlm.nih.gov/pubmed/22323450

MEK • RAF | AKT

PMID:22323450. The principle MAPK pathway activated by the GM CSF receptor is the MEK ERK pathway. Recruitment of mSOS to the SHC GRB2 complex enables mSOS to catalyze RAS activation. Formation of active GTP bound RAS from inactive GDP bound RAS leads to the successive activation of RAF, MEK, and ERK.

PKB (AKT) promotes activation of the MAP kinase signaling cascade, including activation of MAPK1/ERK2, MAPK3/ERK1 and MAPK8/JNK1

1. https://www.ncbi.nlm.nih.gov/pubmed/22323450

ERK • MEK

DUSPG Inactivates MAP kinases. Has a specificity for the ERK family.

1. https://www.ncbi.nlm.nih.gov/pubmed/22323450
https://www.ncbi.nlm.nih.gov/pubmed/9858808

JUN • JNK

MAPK signaling pathway Homo sapiens

1. https://www.genome.jp/dbget-bin/www_bget?hsa04010

JNK • ERK

MAPK signaling pathway Homo sapiens

1. https://www.genome.jp/dbget-bin/www_bget?hsa04010

TAU • ERK

MAPK signaling pathway Homo sapiens

1. https://www.genome.jp/dbget-bin/www_bget?hsa04010

CPLA2 • ERK

MAPK signaling pathway Homo sapiens

1. https://www.genome.jp/dbget-bin/www_bget?hsa04010

FLT3 • PU1

PMID: 24070385. Indeed, Pu.1 directly regulated Flt3 expression on DCs and their precursors in a dose-dependent manner.

1. https://www.ncbi.nlm.nih.gov/pubmed/24070385

GSK3B • !AKT

PMID:22323450. Activity of glycogen synthase kinase 3 (GSK3B), which is negatively regulated by AKT dependent phosphorylation,appears required to avoid human monocyte to macrophage differentiationin monocyte derived DC differentiation cultures.

1. https://www.ncbi.nlm.nih.gov/pubmed/22323450

IKK • CSF2R | AKT

PMID:22323450. GM CSF induced canonical NFkB activation thus appearscrucial to ensure differentiation and survival of DC precursors.

1. https://www.ncbi.nlm.nih.gov/pubmed/22323450

JAK3 • IL4R

PMID: 23124025

PMID: 25159217. The IL4 receptor (IL4R) signals the activation of the Janus kinase 3 (JAK3) STAT6 pathway through its common gamma chain, which leads to the development of immature DCs.

1. https://www.ncbi.nlm.nih.gov/pubmed/25159217
https://www.ncbi.nlm.nih.gov/pubmed/23124025

JAK1 • IL4R

PMID: 27165851. IL4 receptor complexes are deficient of intrinsic kinase activity. Rather, signal transduction is initiated by receptor-associated kinases i.e. a member of JAK family (JAK1, JAK2, JAK3 and TyK2).

1. https://www.ncbi.nlm.nih.gov/pubmed/27165851

SHP1 • USF1:1 & !LnC_DC

PMID: 28465674. LncDC expression is essential for differentiation of human monocytes into dendritic cells. LncDC promotes STAT3 phosphorylation via inhibiting the action of Src homology region 2 domain containing phosphatase 1 (SHP1)

1. https://www.ncbi.nlm.nih.gov/pubmed/28465674

CIITA • STAT5

PMID:22323450. STAT5 promotes expression of MHC class II transactivatorprotein (CIITA), which is essential for transcriptionalactivity of the MHC class II promoter.

1. https://www.ncbi.nlm.nih.gov/pubmed/22323450

ITGAX • IRF4 & PU1 & PRDM1

Sites found using matrix-scan with the regulatory regions of moDC in this study.

PMID: 28338898. PU.1 transactivates the Itgax promoter via direct binding to the cis element on the gene in DCs and through gene regulation of a partner molecule, IRF4, which transactivates the Itgax gene in a synergistic manner with PU.1.

All 11 subpopulations expressed the moDC markers CD1c, CD226, CD48, and CD11c.

moDC marker accoding with Radford et al 2014

1. https://www.ncbi.nlm.nih.gov/pubmed/28338898
https://www.ncbi.nlm.nih.gov/pubmed/29262348
https://pubmed.ncbi.nlm.nih.gov/24513968
https://pubmed.ncbi.nlm.nih.gov/27401672
https://pubmed.ncbi.nlm.nih.gov/29448070

LnC_DC

- PU1 & IRF4 & STAT5

PMID: 28465674
PMID: 24744378
lncDC bound directly to STAT3 in the cytoplasm, which promoted STAT3 phosphorylation on tyrosine705 by preventing STAT3 binding to and dephosphorylation by SHP1.
PU.1 directs lncDC expression in human cDCs

1. https://www.ncbi.nlm.nih.gov/pubmed/28465674
https://www.ncbi.nlm.nih.gov/pubmed/24744378

IL4_gene

- STAT6

DUOX1

- STAT6 & IRF4 & PU1

We observed very specific demethylationin DC differentiation at a CpG site in thegene bodies of DUOX1, an oxidase involved in the antimicrobial mediated response, and the signallingreceptor SLAMF1

1. https://www.ncbi.nlm.nih.gov/pubmed/23124025
https://pubmed.ncbi.nlm.nih.gov/26758199

SLAMF1

- STAT6 & IRF4 & ELK4

We observed very specific demethylationin DC differentiation at a CpG site in thegene bodies of DUOX1, an oxidase involved in the antimicrobial mediated response, and the signallingreceptor SLAMF1

1. https://www.ncbi.nlm.nih.gov/pubmed/23124025
https://pubmed.ncbi.nlm.nih.gov/26758199

MAOA

- STAT6 & NCOR2 & PU1

PMID: 23124025. IL4 can use only the IL4Ralpha,Jak1,Stat3,Stat6 cascade to regulate the expression of some critical inflammatory genes, including ALOX15, monoamine oxidase A (MAOA), and the scavenger receptor CD36.

PMID: 29262348. We found a statistically significant enrichment of genes upregulated in the IL4 signature in MOs GMCSF IL4(0-72h) treated with scrambled siRNAs (Figure6I) whereas genes downregulated in the IL4 signature were enriched in MOs GMCSF IL4(0-72h) treated with antiNCOR2 siRNAs (Figure 6J), establishing NCOR2 as a key regulator for IL4 induced MO differentiation.

1. https://www.ncbi.nlm.nih.gov/pubmed/23124025
https://www.ncbi.nlm.nih.gov/pubmed/29262348

| | |
|---|---|
| HLA_DR<br><br>PMID: 22323450. STAT5 promotes expression of MHC class II transactivatorprotein (CIITA), which is essential for transcriptionalactivity of the MHC class II promoter.<br>PMID: 28465674<br>1. https://www.ncbi.nlm.nih.gov/pubmed/22323450<br>https://www.ncbi.nlm.nih.gov/pubmed/28465674<br>https://pubmed.ncbi.nlm.nih.gov/29262348<br>https://pubmed.ncbi.nlm.nih.gov/29448070 | • STAT3 \| (STAT6 & CIITA) |
| ALOX15<br><br>PMID: 16540365. IL4 can use only the IL4Ralpha,Jak1,Stat3,Stat6 cascade to regulate the expression of some critical inflammatory genes, including ALOX15, monoamine oxidase A (MAOA), and the scavenger receptor CD36.<br>1. pubmed.ncbi.nlm.nih.gov/16540365 | • CREB & STAT6 & STAT3 |
| TIMP3<br>TIMP3 are used to demonstrate the unique regulation of these gene by IL4<br>1. https://pubmed.ncbi.nlm.nih.gov/23124025 | • STAT6 & AP1 & IRF4 |
| DUSP6<br>PMID: 23430108. U0126 treatment significantly reduced the expression of wellknown targets of ERK (FOS, MYC, DUSP6).<br>1. https://www.ncbi.nlm.nih.gov/pubmed/23430108 | • ERK |
| CCL2<br><br>Sites found using matrix-scan with the regulatory regions of moDC in this study.<br><br>PMID: 22328945. ERK upregulated CCL2 expression while impairing the expression of DC maturation markers (RUNX3, ITGB7, IDO1). CCL2, a chemokine constitutively produced by immature MDDCs. monocyte chemoattractant protein 1 (CCL2) have been identified as chemokines/receptors that have an important role in the migration and recruitment of monocytes during the pathogenesis of several inflammatory diseases.<br><br>PMID: 23430108. The CCL2 chemokine directs monocyte/macrophage recruitmentinto tissues under resting and inflamed conditions.<br>1. https://www.ncbi.nlm.nih.gov/pubmed/22328945<br>https://www.ncbi.nlm.nih.gov/pubmed/23430108 | • ERK & STAT5 & STAT3 & FOXO1 |

| | |
|---|---|
| CCL22<br><br>Sites found using matrix-scan with the regulatory regions of moDC in this study.<br><br>PMID: 29262348. We found a statistically significant enrichment of genes upregulated in the IL4 signature in MOs GMCSF IL4(0-72h) treated with scrambled siRNAs (Figure6I) whereas genes downregulated in the IL4 signature were enriched in MOs GMCSF IL4(0-72h) treated with antiNCOR2 siRNAs (Figure 6J), establishing NCOR2 as a key regulator for IL4 induced MO differentiation.<br>1. https://www.ncbi.nlm.nih.gov/pubmed/29262348 | • (AhR & NCOR2 & FOXO1) \| (KLF4 & MAFB) |
| TLR3<br>Sites found using matrix-scan with the regulatory regions of moDC in this study.<br><br>moDC marker accoding with Radford et al 2014<br>1. https://pubmed.ncbi.nlm.nih.gov/24513968 | • IRF4 \| PRDM1 |
| TLR4<br><br>Sites found using matrix-scan with the regulatory regions of moDC in this study.<br><br>moDC marker accoding with Radford et al 2014<br>1. https://pubmed.ncbi.nlm.nih.gov/24513968 | • AP1 \| IRF4 \| PRDM1 \| PU1 |
| TLR6<br><br>Sites found using matrix-scan with the regulatory regions of moDC in this study.<br><br>moDC marker accoding with Radford et al 2014<br>1. https://pubmed.ncbi.nlm.nih.gov/24513968 | • CEBPa \| CEBPb \| STAT6 |
| TLR7<br>Sites found using matrix-scan with the regulatory regions of moDC in this study.<br><br>moDC marker accoding with Radford et al 2014<br>1. https://pubmed.ncbi.nlm.nih.gov/24513968 | • CEBPa \| CEBPb \| IRF4 |
| TLR8<br>Sites found using matrix-scan with the regulatory regions of moDC in this study.<br><br>moDC marker accoding with Radford et al 2014 | • KLF4 \| CEBPa \| STAT6 |

1. https://pubmed.ncbi.nlm.nih.gov/24513968

CD48

- PU1 & IRF4

Sites found using matrix-scan with the regulatory regions of moDC in this study.

All 11 subpopulations expressed the moDC markers CD1c, CD226, CD48, and CD11c.

1. https://www.ncbi.nlm.nih.gov/pubmed/29262348

CD1A

- (BATF3 | CEBPa | CEBPb | CREB) & IRF4 & PU1 & PRDM1 & NCOR2

Sites found using matrix-scan with the regulatory regions of moDC in this study.

PMID:11306493
IL4 signaling upregulates CD1a on cell surface of DC cells.

PMID: 29262348. We found a statistically significant enrichment of genes upregulated in the IL4 signature in MOs GMCSF IL4(0-72h) treated with scrambled siRNAs (Figure6I) whereas genes downregulated in the IL4 signature were enriched in MOs GMCSF IL4(0-72h) treated with antiNCOR2 siRNAs (Figure 6J), establishing NCOR2 as a key regulator for IL4 induced MO differentiation.

1. https://www.ncbi.nlm.nih.gov/pubmed/11306493
https://www.ncbi.nlm.nih.gov/pubmed/29262348
https://pubmed.ncbi.nlm.nih.gov/27401672
https://pubmed.ncbi.nlm.nih.gov/29448070
https://pubmed.ncbi.nlm.nih.gov/17595377

CD1B

- (CEBPa | CEBPb | IRF4) & PRDM1

Sites found using matrix-scan with the regulatory regions of moDC in this study.

Human inflammatory moDC are HLADR CD11c cells that express markers found on classical DC such as CD1c, CD1a, CD1b

1. https://pubmed.ncbi.nlm.nih.gov/29448070

CD1C

- FOXO1 & IRF4 & NR4A1 & PU1 & STAT6

Sites found using matrix-scan with the regulatory regions of moDC in this study.

DC associated (CD1C, ZBTB46) genes.

1. https://www.ncbi.nlm.nih.gov/pubmed/29262348

https://pubmed.ncbi.nlm.nih.gov/29448070

CD40

- AP1

moDC marker accoding with Radford et al 2014

1. https://pubmed.ncbi.nlm.nih.gov/24513968

CD86

- AP1

MOs (GM CSF IL4 ) we identified subsets that either expressed HLA-DR and CD86 or CD1a and FceR1, the former representing a subpopulationwith elevated antigen presenting capacity.

1. https://pubmed.ncbi.nlm.nih.gov/29262348

https://pubmed.ncbi.nlm.nih.gov/27401672

CD83

- STAT6 & NFKB2 & IRF4

Sites found using matrix-scan with the regulatory regions of moDC in this study.

moDCs have been reported to express cel lsurface markers CD80, CD83, CD86,and CD1a in vitro

1. https://pubmed.ncbi.nlm.nih.gov/27401672

CD209

- AP1 & CREB & ELK4 & IRF4 & PU1 & STAT6 & FOXO1

Sites found using matrix-scan with the regulatory regions of moDC in this study.

PMID: 29262348. CD209 was exclusively expressed by MOs GMCS IL4 (0 tp 72h). CD11b didnot discriminate between the cell populations.

moDC marker accoding with Radford et al 2014

1. https://www.ncbi.nlm.nih.gov/pubmed/29262348

https://pubmed.ncbi.nlm.nih.gov/24513968

CD141

- (CEBPa | CREB) & USF1 & ATF1 & IRF4

Gene THBD.

Sites found using matrix-scan with the regulatory regions of moDC in this study.

CD141 phenotypic markers for moDC

1. https://www.genecards.org/cgi-bin/carddisp.pl?gene=THBD&keywords=CD141

https://pubmed.ncbi.nlm.nih.gov/29448070

CD226

- (BATF3 | CEBPa) & FOXO1 & IRF4 & PRDM1 & PU1 &

| | |
|---|---|
| Sites found using matrix-scan with the regulatory regions of moDC in this study.<br><br>All 11 subpopulations expressed the moDC markers CD1c, CD226, CD48, and CD11c.<br>1. https://www.ncbi.nlm.nih.gov/pubmed/29262348 | STAT3 & STAT5 & STAT6 & USF1 |
| DEC205<br><br>Sites found using matrix-scan with the regulatory regions of moDC in this study.<br><br>Gene LY75<br><br>moDC marker accoding with Radford et al 2014<br>1. https://www.genecards.org/cgi-bin/carddisp.pl?gene=LY75&keywords=ly75<br>https://pubmed.ncbi.nlm.nih.gov/24513968 | • FOXO1 & AP1 & PRDM1 |
| DCIR<br>CLEC4A Gene<br><br>Sites found using matrix-scan with the regulatory regions of moDC in this study.<br><br>moDC marker accoding with Radford et al 2014<br>1. https://www.genecards.org/cgi-bin/carddisp.pl?gene=CLEC4A&keywords=DCIR<br>https://pubmed.ncbi.nlm.nih.gov/24513968 | • STAT6 & PU1 |
| Tet2<br>Demethylation is TET2 dependent and is essential for acquiring proper dendritic cell and macrophage identity.<br>1. https://pubmed.ncbi.nlm.nih.gov/26758199 | • PU1 |
| PTPN1<br>PMID: 11694501. The presence of a second aryl phosphate binding site in PTP1B has also been reported by others.<br>1. https://www.ncbi.nlm.nih.gov/pubmed/11694501 | • AhR |
| SOCS<br>PMID: 30578415. However, by far the most important negative regulation occurs at the level of receptor mediated STAT3 activation and is conferred by suppressor of cytokine signalling (SOCS) E3 ubiquitin ligases that enable the degradation of cytokine receptor complexes. SOCS proteins are themselves encoded by STAT target | • STAT3 |

genes and thus provide a transcription dependent negative feedback mechanism.
1. https://www.ncbi.nlm.nih.gov/pubmed/30578415

CD14

- STAT3 & FOXO1 & KLF4 & !STAT5

Sites found using matrix-scan with the regulatory regions of Mo in this study.

PMID: 17762869. We observed an B30 fold induction of the CD14 promoter by KLF4 (Figure 3A).
1. https://www.ncbi.nlm.nih.gov/pubmed/17762869

SELL

- STAT6 & FOXO1 & !PRDM1

PRDM1 is a putative negative regulation, through the sites found using matrix-scan with the regulatory regions of moDC in this study.

Monocyte associated genes (AHR, SELL, CLEC4D)
1. https://pubmed.ncbi.nlm.nih.gov/29262348

CD163

- MAFB & IRF8 & !PRDM1:1

PRDM1 is a putative negative regulation, through the sites found using matrix-scan with the regulatory regions of moDC in this study.

MOs M CSF (macrophages) expressed high amounts of CD163, CD169, and MERTK.
1. https://www.ncbi.nlm.nih.gov/pubmed/29262348

CD206

- MAFB & IRF8 & USF1 & !PRDM1

PRDM1 is a putative negative regulation, through the sites found using matrix-scan with the regulatory regions of moDC in this study.

MOs GM CSF (macrophage) were defined by two subclusters (clusters 7 and 8). Both clusters expressed CD14, CD64, CD68, and CD206.

MR, mannose receptor
1. https://www.ncbi.nlm.nih.gov/pubmed/29262348
https://pubmed.ncbi.nlm.nih.gov/27401672
https://pubmed.ncbi.nlm.nih.gov/10085160

MERTK

- IRF8 & MAFB

Sites found using matrix-scan with the regulatory regions of Mac in this study.

MOs M CSF (macrophages) expressed high amounts of CD163, CD169, and MERTK.
1. https://www.ncbi.nlm.nih.gov/pubmed/29262348

| | |
|---|---|
| CCDC151<br>Sites found using matrix-scan with the regulatory regions of Mac in this study.<br><br>According with Cuevas et al 2017, CCDC151 is an specific gene for macrophages.<br>1. https://pubmed.ncbi.nlm.nih.gov/28093525 | • PU1:1 & AP1 & CEBPb |
| BCL2<br>These results indicate that GM CSF likely induces autophagy by activating JNK and, subsequently, the release of Beclin1 from Bcl2 during monocyte differentiation.<br><br>STAT3 blocks the formation of autophagosomes by driving increased expression of antiautophagic genes Bcl2, Bcl2l1 and Mcl1 and suppression of the proautophagic gene Becn1, which encodes beclin 1.<br>1. https://www.ncbi.nlm.nih.gov/pubmed/30578415 https://www.ncbi.nlm.nih.gov/pubmed/22323450 | • !JNK & STAT3:2 |
| BECN1<br>These results indicate that GM CSF likely induces autophagy by activating JNK and, subsequently, the release of Beclin1 from Bcl2 during monocyte differentiation.<br><br>STAT3 blocks the formation of autophagosomes by driving increased expression of antiautophagic genes Bcl2, Bcl2l1 and Mcl1 and suppression of the proautophagic gene Becn1, which encodes beclin 1.<br>1. https://www.ncbi.nlm.nih.gov/pubmed/30578415 https://www.ncbi.nlm.nih.gov/pubmed/22323450 | • !STAT3 & !BCL2 & JNK |
