## Supplementary material for "Logical modeling of dendritic cells *in vitro* differentiation from human monocytes unravels novel transcriptional regulatory interactions": SuppMaterial

**SupplementatyFigure1**

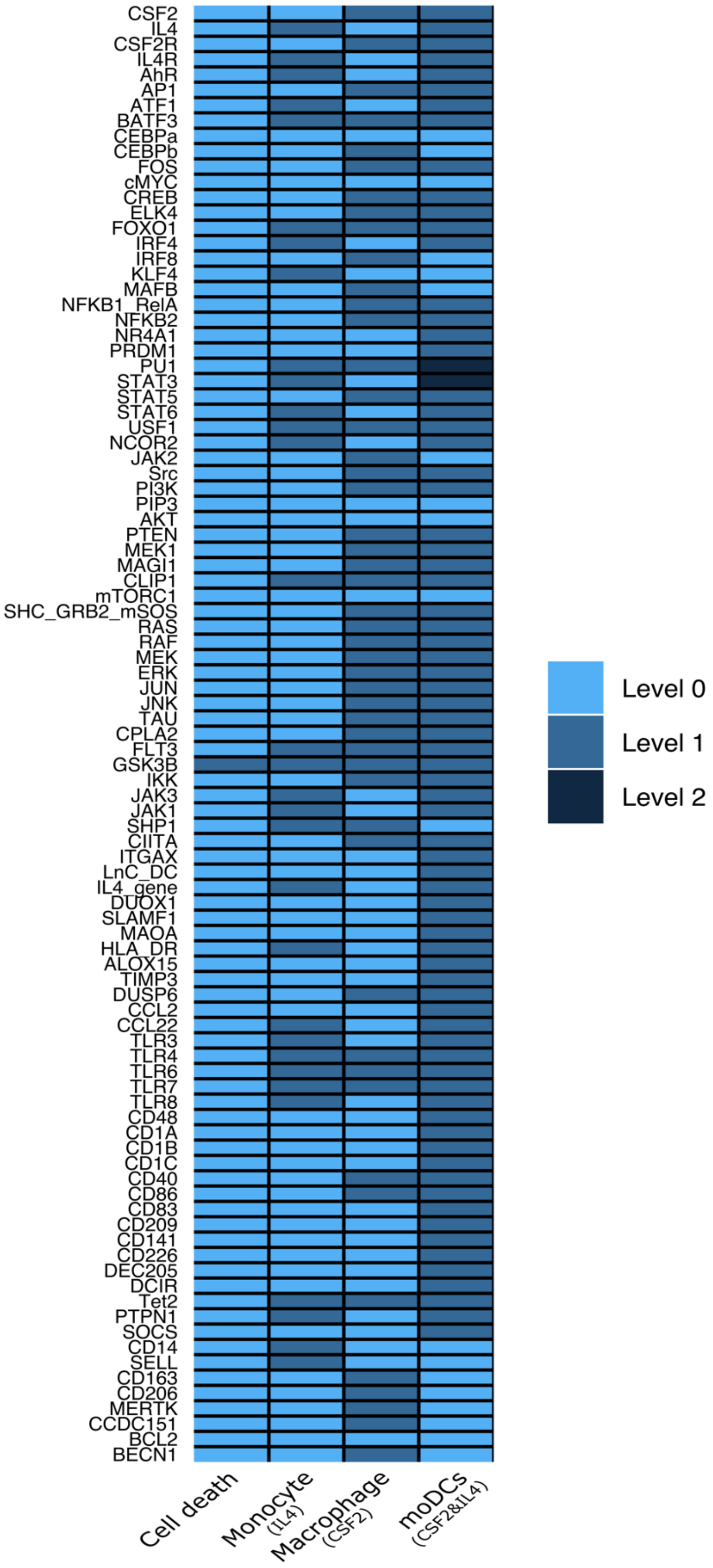

**Supplementary Figure 1.** Stables states of the logical model of monocytes to dendritic cells differentiation *in vitro*. Displays the complete set of nodes, every column corresponds to a specific cell type according with each input. The first column corresponds to the final outcome in the absence of both IL4 and CSF2, *i.e.* cell-death of the monocytes. The second column corresponds to the stimulation of monocytes by IL4. The third column corresponds to the macrophage outcome, in the presence of the sole CSF2. Finally, the fourth column corresponds to moDCs commitment, in the presence of both IL4 and CSF2, where STAT3 reaches the level 2 in the presence of the long non-coding RNA LnC-DC, and PU.1 reaches the level 2, which is required to turn-off MAFB during moDCs commitment.

| Sample ID | TechnicalReplicate | #RawReads | #MappedReads | %Mapped | #UniquelyMappedReads | %UniquelyMapped | PBC | NRF | NSC | RSC |
|---|---|---|---|---|---|---|---|---|---|---|
| EGAR00001085103 | none | 36270988 | 34556576 | 95.27 | 34429873 | 94.92 | 0.62 | 0.61 | 1.213468 | 4.973275 |
| EGAR00001088389 | none | 59464947 | 58507187 | 98.39 | 58195238 | 97.86 | 0.54 | 0.5 | 1.193545 | 3.120265 |
| EGAR00001402942 | none | 61298322 | 58636670 | 95.66 | 58494233 | 95.43 | 0.72 | 0.72 | 1.12851 | 2.988048 |
| EGAR00001402945 | none | 53792767 | 50450609 | 93.79 | 50276262 | 93.46 | 0.72 | 0.72 | 1.096921 | 2.880922 |
| EGAR00001096292 | none | 43643430 | 43124385 | 98.81 | 42914833 | 98.33 | 0.62 | 0.6 | 1.144857 | 2.705355 |
| EGAR00001088409 | none | 34883406 | 34269455 | 98.24 | 34089818 | 97.73 | 0.75 | 0.75 | 1.122595 | 2.300364 |
| EGAR00001305774 | none | 62396291 | 52634872 | 84.36 | 52526559 | 84.18 | 0.71 | 0.71 | 1.673187 | 1.496055 |
| EGAR00001402937 | none | 48107687 | 46457099 | 96.57 | 46352205 | 96.35 | 0.71 | 0.71 | 1.638107 | 1.480712 |
| EGAR00001402975 | none | 53756793 | 52000602 | 96.73 | 51916823 | 96.58 | 0.7 | 0.68 | 2.603013 | 1.158121 |
| EGAR00001402944 | none | 46489075 | 45459055 | 97.78 | 45336634 | 97.52 | 0.94 | 0.94 | 1.06705 | 1.153417 |
| EGAR00001080465 | none | 58927868 | 57311046 | 97.26 | 56365411 | 95.65 | 0.59 | 0.52 | 1.155807 | 1.144716 |
| EGAR00001305771 | none | 50554614 | 49043003 | 97.01 | 48942079 | 96.81 | 0.94 | 0.93 | 1.169745 | 1.077844 |
| EGAR00001085095 | none | 26253843 | 25776307 | 98.18 | 25671563 | 97.78 | 0.72 | 0.72 | 1.051644 | 0.8222461 |
| EGAR00001402946 | none | 49647095 | 47554723 | 95.79 | 46817138 | 94.3 | 0.89 | 0.84 | 1.052285 | 0.4493812 |
| EGAR00001320582 | none | 35464413 | 34464338 | 97.18 | 34301686 | 96.72 | 0.98 | 0.97 | 1.012696 | 0.3588252 |
| EGAR00001402965 | none | 34302091 | 33342092 | 97.2 | 33187744 | 96.75 | 0.98 | 0.98 | 1.008811 | 0.2364044 |
| EGAR00001402963 | none | 33958907 | 33557992 | 98.82 | 33394119 | 98.34 | 0.98 | 0.98 | 1.008083 | 0.2085872 |
| EGAR00001228104 | none | 25945154 | 25256088 | 97.34 | 25189608 | 97.09 | 0.88 | 0.88 | 1.176624 | 1.884106 |
| EGAR00001080466 | none | 44148235 | 40264959 | 91.2 | 40082521 | 90.79 | 0.58 | 0.56 | 1.718232 | 1.886858 |
| EGAR00001100410 | none | 44271422 | 43530938 | 98.33 | 43305088 | 97.82 | 0.88 | 0.87 | 1.047842 | 1.455561 |
| EGAR00001231891 | none | 36632043 | 36058094 | 98.43 | 35867916 | 97.91 | 0.73 | 0.73 | 1.140675 | 2.899751 |
| EGAR00001132113 | none | 40076490 | 27355246 | 68.26 | 27213222 | 67.9 | 0.84 | 0.83 | 1.130482 | 2.327079 |
| EGAR00001100397 | none | 44848605 | 43388220 | 96.74 | 43164756 | 96.25 | 0.87 | 0.87 | 1.103035 | 1.438625 |
| EGAR00001143021 | none | 28239437 | 24486109 | 86.71 | 24400038 | 86.4 | 0.79 | 0.78 | 1.154481 | 3.878428 |
| EGAR00001132117 | none | 56770007 | 50780835 | 89.45 | 50022736 | 88.11 | 0.76 | 0.73 | 1.096564 | 0.9502099 |
| EGAR00001132116 | none | 52456200 | 41300163 | 78.73 | 41112241 | 78.37 | 0.76 | 0.75 | 1.120459 | 2.763786 |

**Supplementary table 1.** QC table results after alignment. QC results of the samples used for ChIP-seq data analysis.

| TF | JASPAR ID |
|---|---|
| AHR | MA0006.1 |
| AP1 | MA0940.1 |
| ATF1 | MA0604.1 |
| BATF3 | MA0835.1 |
| CEBPA | MA0102.3 |
| CEBPB | MA0466.2 |
| CREB1 | MA0018.3 |
| ELK4 | MA0076.1 |
| FOXO1 | MA0480.1 |
| IRF4 | MA1419.1 |
| IRF8 | MA0652.1 |
| KLF4 | MA0039.2 |
| MAFB | MA0117.2 |
| NFKB1 | MA0105.4 |
| NFKB2 | MA0778.1 |
| NR4A1 | MA1112.1 |
| SPI1 | MA0080.4 |
| STAT3 | MA0144.2 |
| STAT5 | MA0519.1 |
| STAT6 | MA0520.1 |
| USF1 | MA0093.3 |
| PRDM1 | MA0508.3 |

**Supplementary table 2.** Transcription factors JASPAR ID. List of the transcription factors and their corresponding ID from the JASPAR database that were used in this study to find transcription factors binding sites.
